# Singlet-states-mediated highly efficient two-photon fluorescence in organic dyes

**DOI:** 10.64898/2026.06.04.729720

**Authors:** Shunxin Wang, Xiaoxiao Fan, Xiaofei Miao, Derong Fan, Xiaolong Liu, Zhe Feng, Wenbo Hu, Jun Qian

## Abstract

This study reports a brand-new continuous-wave-excited (CW-excited) two-photon fluorescence emission mechanism in indocyanine green (ICG), an organic fluorescent dye widely used in clinical practice. This mechanism is based on the excited state absorption (ESA) process of the first singlet excited state. Intramolecular electrons sequentially absorb two photons to reach a high-energy singlet excited state, followed by direct radiative transition to the ground state to generate fluorescence. The entire process is exclusively mediated by singlet states. We further summarize the essential requirements for organic dyes to realize this luminescence mechanism. First, the dye must possess at least two well-separated singlet excited states with distinct energies, corresponding to two absorption peaks at different wavelengths in the absorption spectrum. The wavelength of the high-energy singlet excited state is about half that of the first singlet excited state. Second, the peak in the absorption spectrum corresponding to the transition from the ground state to the first singlet excited state has a sufficiently large molar extinction coefficient. Third, the first singlet excited state exhibits the capability of ESA. Fourth, electrons at the high-energy singlet excited state can directly transit to the ground state and emit fluorescence. We validated this mechanism in a variety of organic dyes satisfying the above conditions, confirming its universality. Using CW laser as the excitation source, we achieved two-photon fluorescence imaging of mouse cerebral blood vessels at a depth of 400 μm, which clearly resolves three-dimensional vascular networks with high resolution. We also performed two-photon fluorescence imaging on human gastric cancer tissue samples at a depth of around 150 μm, which provides a low-cost strategy for clinicians to rapidly acquire high-contrast tumor tissue images.

## Introduction

Two-photon fluorescence microscopy serves as a vital tool for fundamental research and translational applications in biology, clinical medicine and related fields^1^. Benefiting from its inherent optical sectioning capability and high signal collection efficiency^2^, it enables high-resolution three-dimensional optical sectioning imaging of cells, highly scattering biological tissues and organs^3–5^. However, conventional two-photon fluorescence microscopy relies on femtosecond pulsed lasers with high peak power as excitation sources. Such systems are expensive and pose safety hazards. Additionally, the excessively high peak power readily induces photobleaching of fluorescent dyes.

Here, we propose a novel two-photon fluorescence emission mechanism in organic dye systems, which can be excited by continuous-wave (CW) lasers. This mechanism is fully realized via transitions between singlet states of intramolecular electrons. Under CW laser excitation, electrons in the ground state first absorb one photon to reach the first singlet excited state. They then absorb another photon with the same energy through excited-state absorption (ESA) and reach the high-energy singlet excited state. Finally, electrons undergo a direct radiative transition from the high-energy singlet excited state to the ground state, generating fluorescence. The involvement of the intermediate singlet excited state greatly boosts two-photon absorption efficiency, and thus enhances two-photon fluorescence emission efficiency. This mechanism is applicable to small organic molecules and organic nanoparticles. Typical examples include indocyanine green (ICG), an FDA-approved near-infrared (NIR) fluorescent dye with more than 60 years of clinical application, and OTL38, a recently FDA-approved dye capable of targeting ovarian cancer tumors. On this basis, we built a two-photon fluorescence microscopic system excited by CW laser. High-resolution optical sectioning imaging was successfully performed on cerebral blood vessels of living mice and human gastric cancer tissues.

## Result

### Spectral properties and energy level diagram of ICG

Earlier attempts have adopted CW lasers to excite two-photon fluorescence of organic dyes, yet the efficiency remained low. Both theoretical and experimental results demonstrate that this approach requires extremely high CW laser power, because the two-photon absorption relies on virtual energy states^6,7^. By contrast, the intermediate singlet excited state can significantly improve the two-photon absorption efficiency, thereby enhancing two-photon fluorescence emission^8,9^.

Fig. 1a shows the steady-state absorption spectrum of ICG. Besides the major absorption peak at 795 nm, a secondary absorption band appears around 400 nm, which is presumed to correspond to high-energy excited states. To verify this hypothesis, we calculated the simulated absorption spectrum of ICG via Time-Dependent Density-Functional Theory (TD-DFT) (Fig. 1b). In addition to the singlet excited state S_1_ corresponding to the main absorption peak, the secondary band is assigned to multiple singlet excited states, collectively denoted as S_n_. To clarify the fluorescence emission pathways from S_n_, we irradiated ICG solution with CW laser at 440 nm. The fluorescence spectrum is presented in Fig. 1c, covering visible and NIR regions. The visible fluorescence with a peak at 555 nm originates from direct radiative transition from S_n_ to the ground state S_0_. The NIR fluorescence peaks at 822 nm, generated by radiative transition from S_1_ to S_0_ after electrons undergo internal conversion (IC) from S_n_ to S_1_. The molar extinction coefficient of the main absorption peak reaches approximately 3.75×10^5^ L·mol^-1^·cm^-1^, enabling rapid accumulation of sufficient electrons in S_1_. Moreover, the wavelength of the main absorption peak is roughly twice that of the other peak. Hence, we infer that electrons in S_0_ can sequentially absorb two photons (e.g., 800 nm) and reach the high-energy state S_n_ via a two-step process. To investigate this, the transient absorption spectroscopy of ICG was measured using a femtosecond excitation source centered at 800 nm (Fig. 1d). Obvious ESA was observed around 800 nm.

**Fig. 1.**
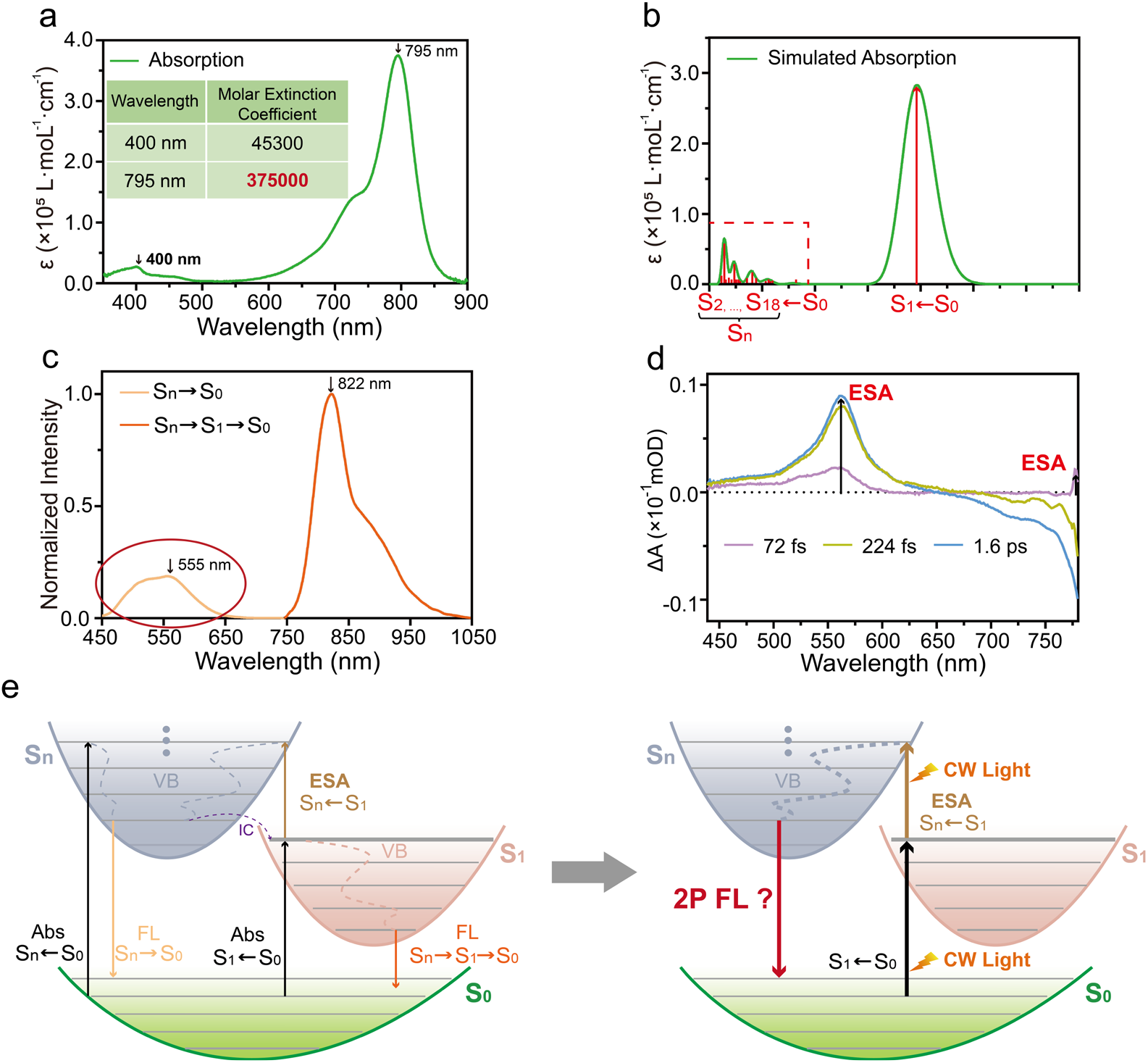
Spectral properties and the complete Jablonski energy level diagram of ICG. **a**. Steady-state absorption spectrum of ICG in DMSO solution. **b**. TD-DFT simulated absorption spectrum of ICG. **c**. Fluorescence spectrum of ICG under 440 nm CW excitation. **d**. Transient absorption spectrum of ICG under 800 nm femtosecond laser excitation. **e**. Complete Jablonski energy level diagram and hypothesis of CW-excited two-photon fluorescence of ICG.

According to the above findings, we established the complete Jablonski energy diagram of ICG (Fig. 1e). When illuminated by photons with a wavelength around 800 nm, electrons in the S_0_ state can first absorb a photon and transition to the S_1_ state. From there, the electrons can either directly return to S_0_ via radiative transition, emitting NIR fluorescence, or undergo the ESA process by absorbing another photon of the same energy to reach the S_n_ state. Once at the S_n_ state, the electrons can either internally convert back to S_1_ and subsequently return to S_0_ via radiative transition, emitting NIR fluorescence, or directly undergo a radiative transition from S_n_ to S_0_, emitting visible fluorescence. It should be noted that the transition of electrons from S_0_ to S_1_ corresponds to a large molar extinction coefficient in the absorption spectrum, which facilitates the rapid population of a sufficient number of electrons in the S_1_ state within a short time, thereby promoting the ESA process with high efficiency. Accordingly, we raise a question: under CW excitation near 800 nm, can electrons absorb two photons successively via S_1_ to S_n_, and then emit two-photon fluorescence through radiative transition back to S_0_ directly?

### CW-excited two-photon fluorescence of ICG

The CW laser operating at 785 nm was employed for the excitation of ICG. The fluorescence spectrum of ICG were measured in two spectral ranges: below 700 nm and above 800 nm (Fig. 2a). The results show that ICG emits fluorescence in the visible region besides the well-known NIR fluorescence, with a peak at about 564 nm within the band of 500-650 nm. This peak is close to the visible emission peak under 440 nm CW excitation. Subsequently, the CW laser was coupled into a commercial two-photon laser scanning fluorescence microscope for imaging. The microscope was equipped with two detection channels: a green channel covering 500-575 nm and a red channel covering 575-650 nm. Distinct fluorescence signals of ICG were detected in both channels. Next, a cuvette containing ICG solution was placed under the objective lens and excited by focused 785 nm CW laser. Fluorescence spots were captured laterally with a camera. When the detection range was set to 500-650 nm, the fluorescence spot appeared as a dot (Fig. 2b), whereas a line was observed when the detection range was above 900 nm (Fig. 2c), indicating that the fluorescence in the 500-650 nm band exhibits spatially confined characteristics. The relationship between fluorescence intensity and excitation power was measured (Fig. 2d), and the fitted slopes were consistently close to 2, indicating that the fluorescence within the band of 500-650 nm under 785 nm CW excitation is a two-photon emission process. As the CW laser power gradually increased, the slope of the fitted line decreased markedly (Fig. 2e), suggesting the presence of excited-state saturation, which supported the involvement of the intermediate singlet excited state in this two-photon fluorescence process. In contrast, the power dependence of fluorescence within the band of 650-700 nm was linear over the same excitation power range (Fig. 2f), indicating that this spectral component in Fig. 2a arose from the left tail of the NIR one-photon fluorescence. In addition, fluorescence lifetimes of the emission within the band of 500-650 nm were measured using 440 nm and 790 nm excitation sources (Fig. 2g, h). The values were 221 ± 12 ps and 219 ± 15 ps, respectively, which were comparable and suggested that the two-photon fluorescence originated from direct transitions from the S_n_ to S_0_. The complete process was consistent with the scheme proposed in Fig. 1e. Finally, the photostability of the two-photon fluorescence of ICG was evaluated under continuous 785 nm CW laser irradiation for 18 min, during which ICG maintained a remarkably bright two-photon fluorescence signal (Fig. 2i), supporting its applicability for in vivo two-photon fluorescence imaging.

**Fig. 2.**
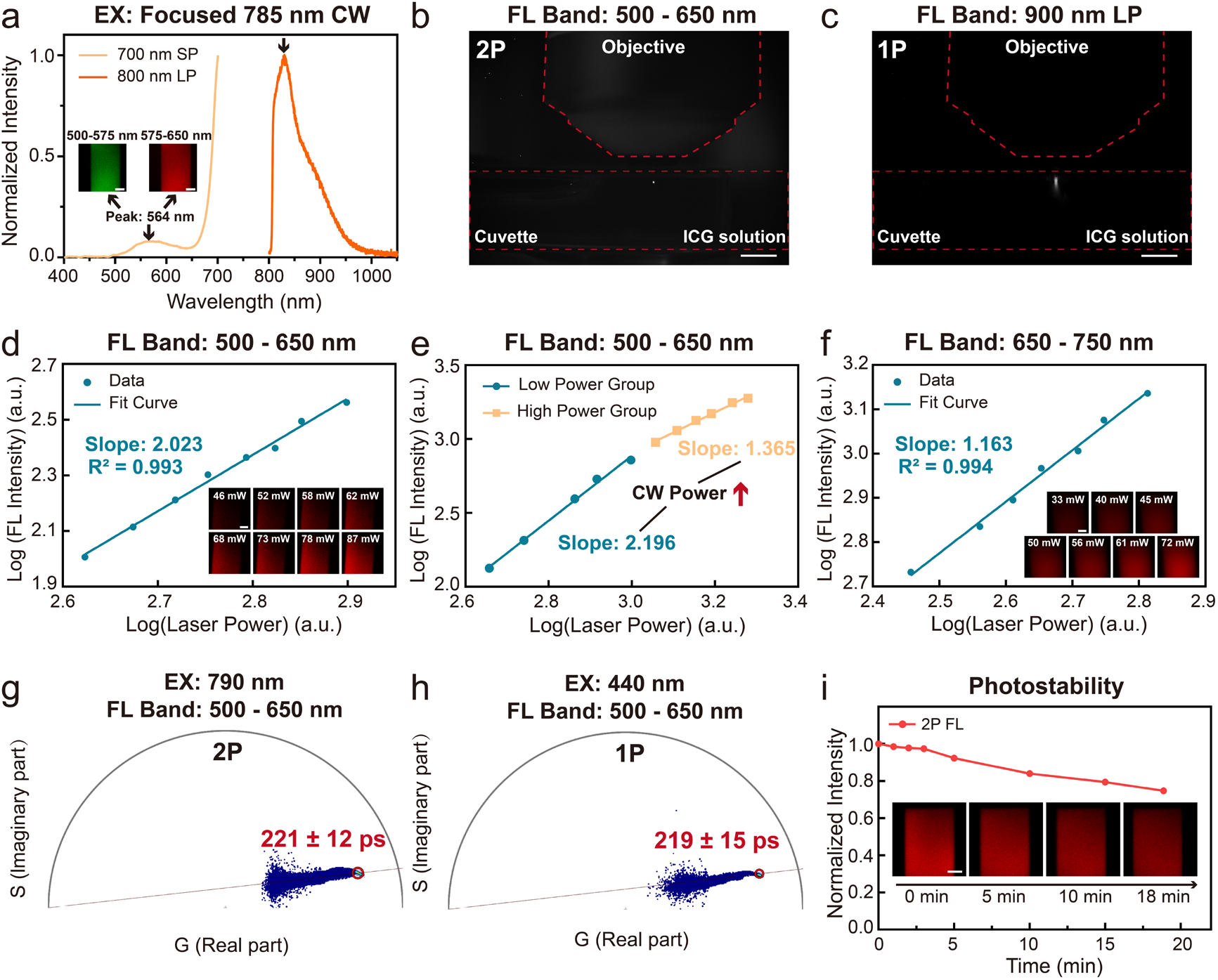
Validation of two-photon fluorescence of ICG under CW excitation. **a**. Fluorescence spectrum of ICG under 785 nm CW excitation. **b-c**. Fluorescence spot images of ICG solution in different wavelength bands under focused 785 nm CW excitation: b. 500–650 nm band, exposure time: 90 ms. c. 900 nm long-pass band, exposure time: 1 ms. **d**. Relationship between fluorescence intensity in the 500-650 nm band of ICG and 785 nm CW excitation power. **e**. Relationship between fluorescence intensity in the 500-650 nm band of ICG and excitation power, two segments corresponding to the relatively low power range and high power range respectively. **f**. Relationship between fluorescence intensity in the 650–750 nm band of ICG and 785 nm CW excitation power. **g**. Two-photon fluorescence lifetime of ICG under 790 nm excitation. **h**. One-photon fluorescence lifetime of ICG in the same band under 440 nm excitation. **i**. Photostability of two-photon fluorescence under 785 nm CW laser excitation. Scale bars: 100 μm for the fluorescence images in Fig. 2a, d, f, i; 5 mm for Fig. 2b, c.

### Generalization of the proposed mechanism

Based on the above theoretical basis and the intrinsic characteristics of ICG, we summarize that organic dyes must simultaneously meet the following conditions to achieve similar CW-excited two-photon fluorescence (Fig. 3a). First, they should possess two sufficiently separated absorption bands, with the long-wavelength absorption peak approximately twice that of the short-wavelength absorption peak. Second, the molar extinction coefficient of the long-wavelength absorption band should be sufficiently large, on the order of ~10^5^, which facilitates the population of a sufficient number of electrons in the intermediate singlet state within a short time. Third, the intermediate singlet excited state should exhibit ESA to reach a higher-energy singlet excited state, with the first two conditions serving as the basis for this requirement. Fourth, electrons should be able to directly transition from the high-energy singlet excited state to the ground state and emit fluorescence.

**Fig. 3.**
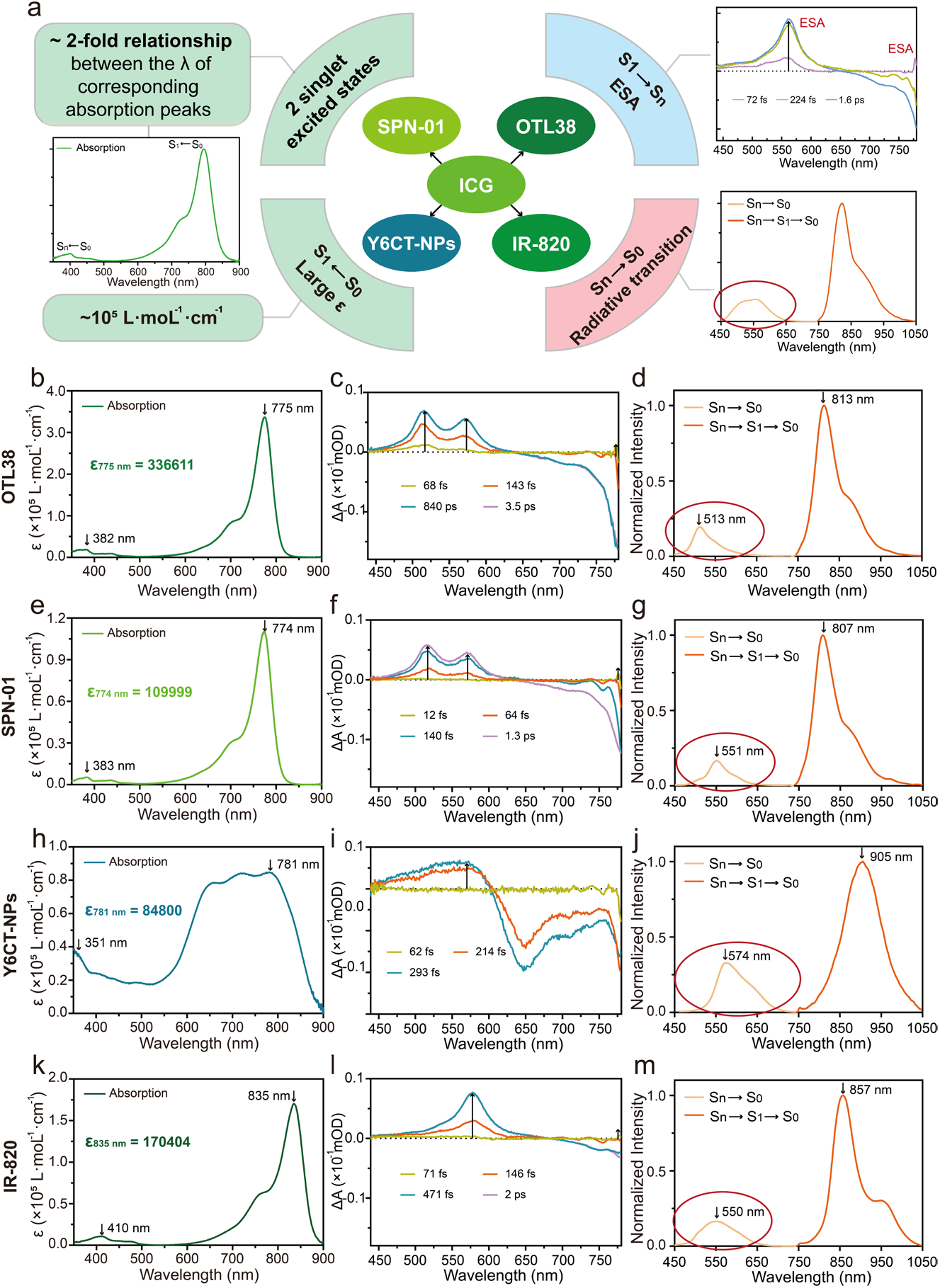
Generalization of the proposed mechanism. **a**. Schematic diagram of the requirements for organic dyes to adopt this mechanism. **b, e, h, k**. Steady-state absorption spectra of OTL38, SPN-01, Y6CT-NPs in aqueous solution and IR-820 in DMSO solution, respectively. **c, f, i, l**. Femtosecond transient absorption spectra of the above samples upon 800 nm excitation. **d, g, j, m**. Fluorescence spectra of the above samples under 440 nm CW excitation.

Based on these criteria, we identified several organic dyes that meet the above conditions. The first is OTL38, an FDA-approved NIR organic fluorescent small molecule with ovarian cancer tumor-targeting ability. Its aqueous solution exhibits two well-separated absorption bands with peak wavelengths at 382 nm and 775 nm (Fig. 3b), and possesses a large molar extinction coefficient at 775 nm, approximately 3.3 × 10^5^ L·mol^-1^·cm^−1^. Transient absorption spectrum confirmed the presence of ESA around 800 nm (Fig. 3c). Meanwhile, the aqueous solution of OTL38 exhibits visible fluorescence emission with a peak wavelength of 513 nm under 440 nm CW excitation (Fig. 3d). The second is SPN-01, a NIR organic fluorescent molecule. It is conjugated with an arginine-glycine-aspartic acid (RGD) peptide. Because the RGD peptide can bind to integrins overexpressed on tumor vascular endothelial cells, this molecule possesses tumor-targeting capability. The aqueous solution of SPN-01 also displays two well-separated absorption bands with peak wavelengths at 383 nm and 774 nm (Fig. 3e), and a large molar extinction coefficient at 774 nm, approximately 1.1 × 10^5^ L·mol^−1^·cm^−1^. Transient absorption spectrum confirmed the presence of ESA around 800 nm (Fig. 3f). Under 440 nm CW excitation, SPN-01 emits visible fluorescence with a peak wavelength of 551 nm (Fig. 3g). Both OTL38 and SPN-01 described above are organic single molecules. Next, we found that an organic nanoparticle, Y6CT-NPs, also meets the above conditions. The aqueous solution of Y6CT-NPs exhibits a broad absorption spectrum covering the visible and NIR regions, but can still be roughly divided into two absorption bands with peak wavelengths at 351 nm and 781 nm respectively (Fig. 3h). It has a relatively large molar extinction coefficient at 781 nm, approximately 8.5 × 10^4^ L·mol^−1^·cm^−1^. Transient absorption spectrum confirmed the presence of ESA (Fig. 3i). Similarly, it emits visible fluorescence with a peak wavelength of about 574 nm under 440 nm CW excitation (Fig. 3j). In addition, IR-820, an analogue of ICG, also satisfies the above conditions (Fig. 3k-m).

Subsequently, we sequentially verified the CW-excited two-photon fluorescence of the above organic molecules and nanoparticles. The corresponding results are presented in Fig. 4. Under 785 nm CW excitation, all four organic dyes emitted fluorescence in the visible region (Fig. 4a, d, g, j). For each dye, we measured the power dependence of fluorescence intensity within the 500-650 nm range in three independent tests. The fitted slopes remained consistently around 2, confirming the two-photon fluorescence emission. As the excitation power increased gradually, the fitted slopes all declined (Fig. 4c, f, i, l), further demonstrating that intermediate singlet excited states are involved in the CW-excited two-photon fluorescence process of these dyes.

**Fig. 4.**
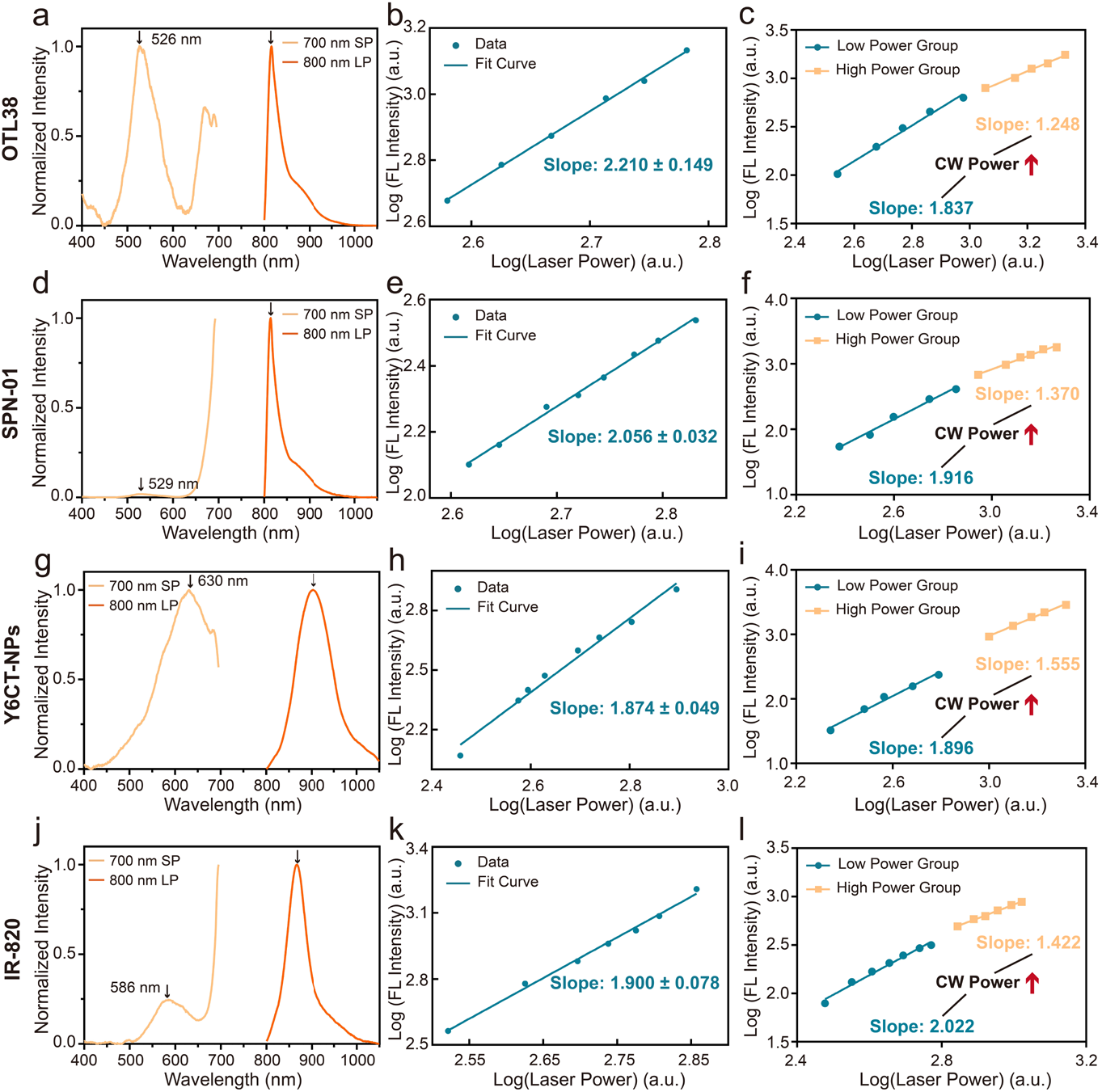
Validation of CW-excited two-photon fluorescence of different organic dyes. **a, d, g, j.** Fluorescence spectra of different organic dye solutions under 785 nm CW excitation. **b, e, h, k**. Relationships between fluorescence intensity in the 500-650 nm band and 785 nm CW excitation power of different organic dye solutions. **c, f, i, l**. Relationships between fluorescence intensity in the 500–650 nm band and 785 nm CW excitation power of different organic dye solutions, two segments corresponding to the relatively low power range and high power range respectively.

In addition, two-photon and one-photon fluorescence lifetime measurements in the same visible band (500–650 nm) were performed for the above organic dyes, as shown in Fig. 5. For each dye, the one-photon fluorescence lifetime under 440 nm excitation was close to the two-photon fluorescence lifetime under 790 nm excitation, indicating that the two-photon fluorescence originated from a high-energy singlet state. Furthermore, under continuous 785 nm CW irradiation, all these organic dyes exhibited good photostability.

**Fig. 5.**
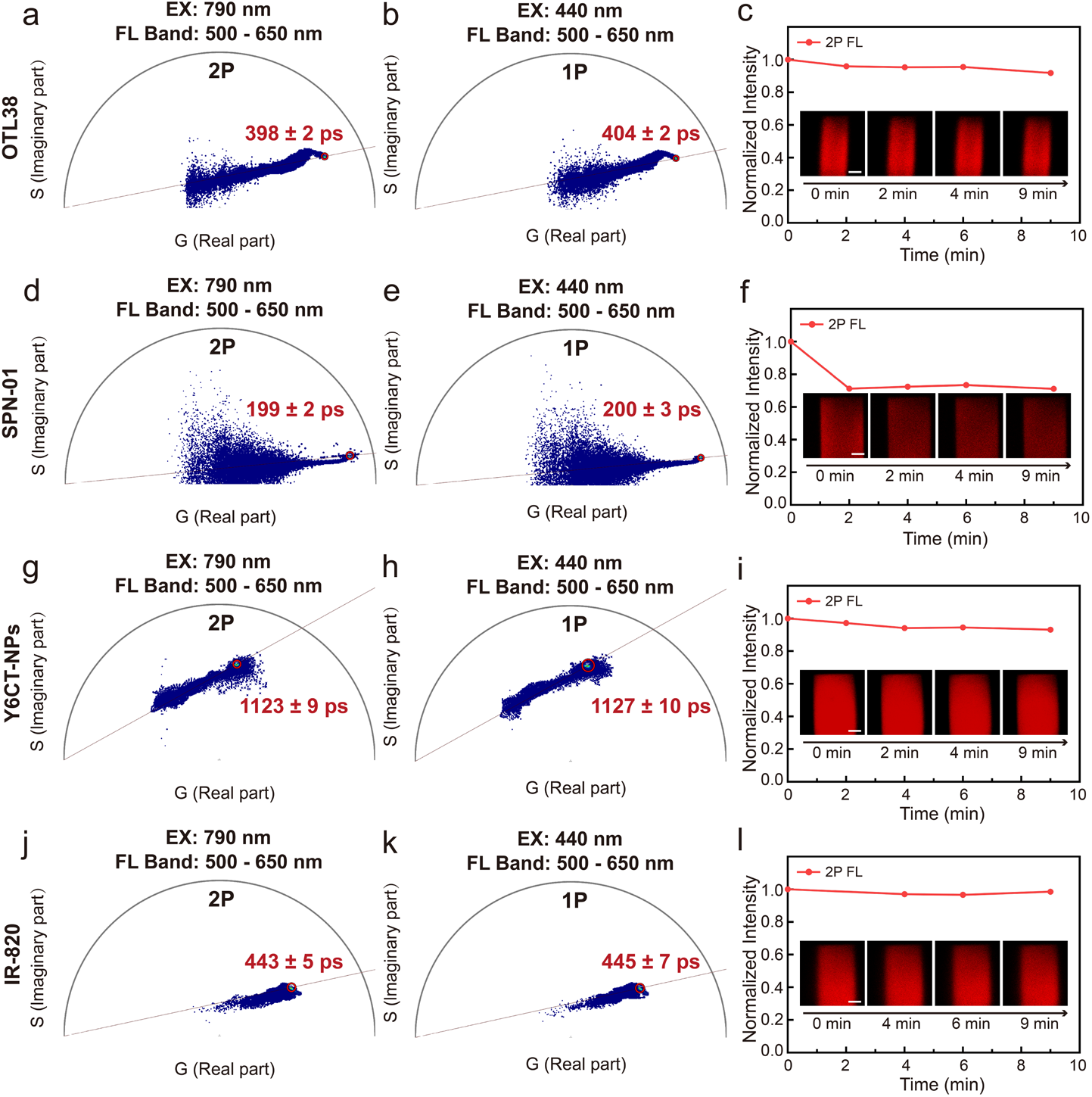
Two-photon fluorescence lifetimes and photostability of different organic dyes. **a, d, g, j**. Two-photon fluorescence lifetimes of different organic dye solutions under 790 nm laser excitation. **b, e, h, k**. one-photon fluorescence lifetimes of different organic dye solutions under 440 nm laser excitation. **c, f, i, l**. two-photon fluorescence photostability of different organic dye solutions under 785 nm CW excitation. Scale bars: 100 μm.

Collectively, we have demonstrated ESA-based CW-excited two-photon fluorescence emission in three different systems of organic fluorophores: small molecules, nanoparticles, and small molecules conjugated with peptides. These findings support the general applicability of the proposed mechanism.

### CW-excited two-photon fluorescence imaging of mouse cerebral vasculature

In this section, taking mouse cerebral vasculature imaging as a representative example, we validate the capability of this mechanism for high-resolution three-dimensional optical sectioning imaging in highly scattering biological media. The setup of the CW-excited two-photon fluorescence microscope is shown in Fig. 6a. The system is based on the frame of a commercial two-photon scanning microscope, incorporating a 785 nm CW laser. The CW laser beam passes through a two-dimensional galvanometric scanning mirror, a scan lens, and a tube lens, and is finally focused by a high numerical aperture (NA) objective for point scanning. The generated two-photon fluorescence is collected by the same objective, reflected by a dichroic mirror to separate it from the excitation light, and finally detected by a photomultiplier tube (PMT).

**Fig. 6.**
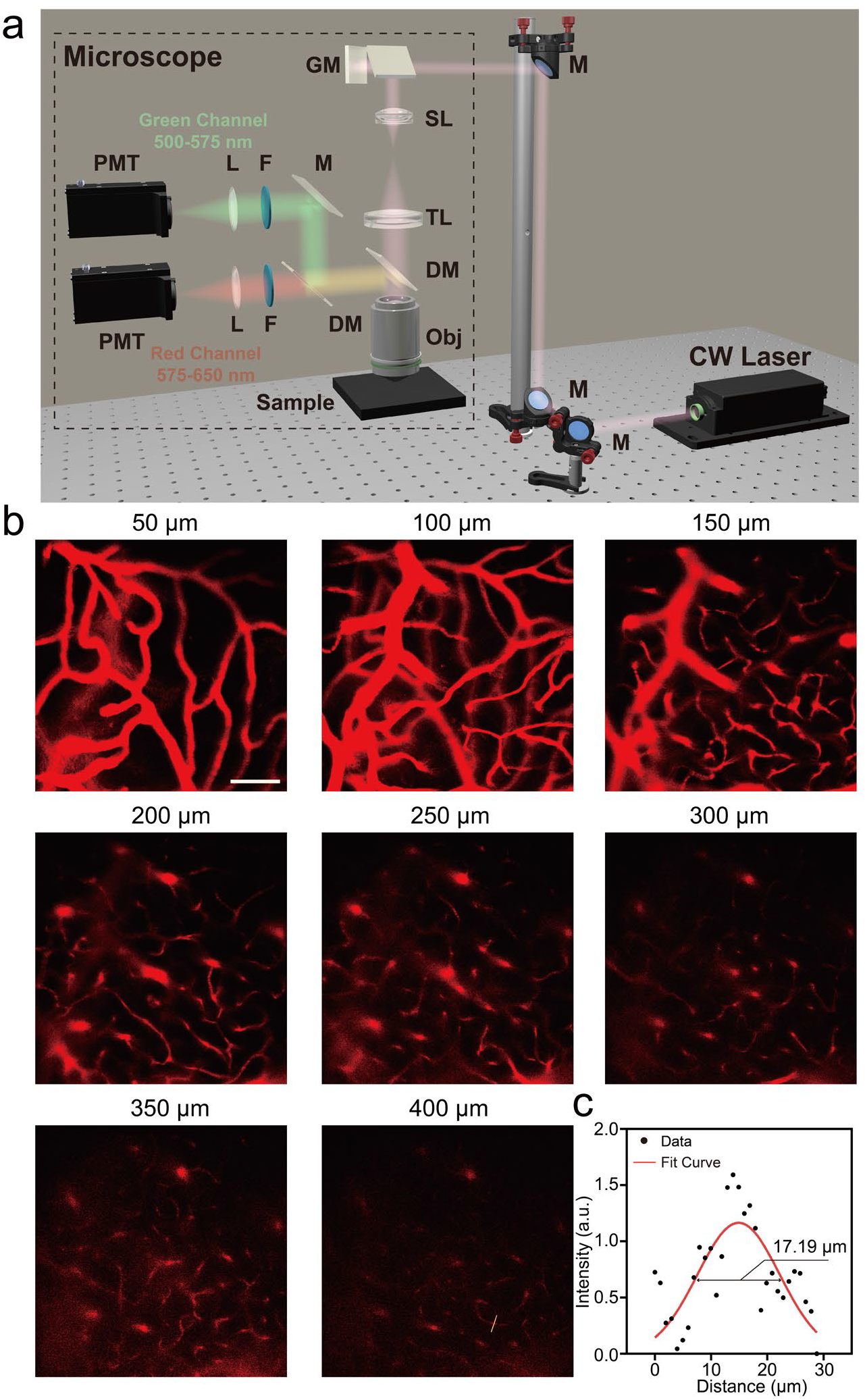
CW-excited two-photon fluorescence imaging of mouse cerebral vasculature labeled with ICG. **a**. Schematic of the CW-excited two-photon fluorescence microscope. **b**. Two-photon fluorescence images of mouse cerebral vasculature at different depth planes. Pixel number: 512 × 512, pixel dwell time: 2 μs, PMT bias voltage: 650 V for superficial layers and 750 V for the deepest layer. Scale bar: 100 μm. **c**. Fluorescence intensity profile along the yellow line in the image at a depth of 400 μm, together with the Gaussian fitting result.

After craniotomy on the mouse, ICG was intravenously injected to label the cerebral vasculature. Under 785 nm CW excitation, fluorescence in the 500–650 nm band was selected for detection. Two-photon fluorescence images of mouse cerebral vasculature at different depths were obtained (Fig. 6b), revealing a rich vascular network with clearly discernible vessels. The maximum imaging depth was approximately 400 μm. These results demonstrate that CW-excited two-photon fluorescence possesses excellent optical sectioning capability. Compared with femtosecond lasers, CW lasers are inexpensive and easy to operate. Taken together, this mechanism provides a lower-cost and simpler solution for high-resolution three-dimensional imaging in highly scattering media.

### CW-excited two-photon fluorescence imaging of human gastric cancer tissue

In this section, CW-excited two-photon fluorescence imaging was performed on clinical human gastric cancer tissues. Paired fresh gastric cancer tissues and adjacent normal tissues were obtained from the operating room. Both types of tissue samples were rapidly immersed in the aqueous solution of OTL38 and incubated for 30 min, followed by three washes with PBS buffer within 30 min. Without sectioning, the samples were directly placed under the objective lens for imaging. The CW-excited two-photon fluorescence images of the tumor samples incubated with OTL38 solution at different depths are shown in Fig. 6b, achieving optical sectioning imaging down to a depth of 150 μm. Representative punctate structures, corresponding to folate receptor clusters on the tumor cell membrane, were observed at various depths. In contrast, the adjacent normal tissue exhibited almost no fluorescence signal (Fig. 6c). Second Near-infrared Window (NIR-II, 900-1880 nm) macroscopic fluorescence imaging revealed that the washed tumor tissue still displayed bright NIR-II fluorescence, whereas the adjacent normal tissue showed negligible signal (Fig. 6d). These results were consistent with those of H&E staining (Fig. 6e,f).

In summary, CW-excited two-photon fluorescence enables optical sectioning imaging of thick tumor tissues with higher scattering to a certain depth without the need for frozen sectioning, and effectively distinguishes tumor tissues from adjacent normal tissues. This approach can assist physicians in rapidly acquiring high-contrast tissue images, substantially reducing the time cost of diagnosis. The low-cost, flexible, easy-to-maintain, and safer CW light source also facilitates its deployment and use in various surgical departments.

**Fig. 7.**
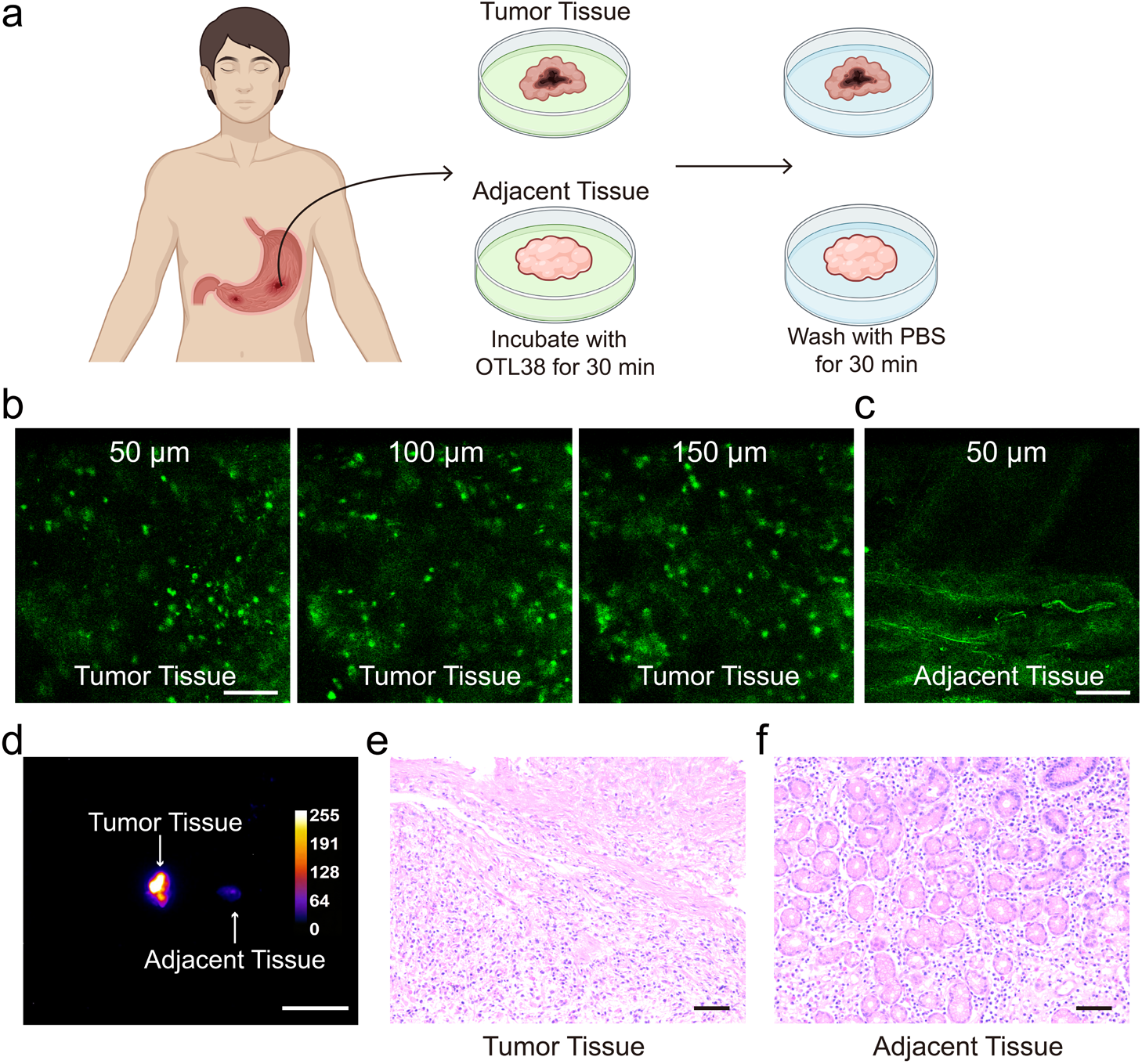
CW-excited two-photon fluorescence imaging of human gastric cancer tissue. **a**. Schematic diagram of the tissue processing procedure. **b**. Two-photon fluorescence images of human tissue at different depths under 785 nm CW excitation. Pixel number: 512 × 512, pixel dwell time: 2 μs, PMT bias voltage: 660 V for the superficial layer and 800 V for the deepest layer. **c**, Two-photon fluorescence image of normal tissue at a depth of 50 μm. Pixel number: 512 × 512, pixel dwell time: 2 μs, PMT bias voltage: 800 V. **d**. NIR-II fluorescence images of tumor and adjacent normal tissues. Excitation power density: 10 mW/cm^2^. Exposure time: 30 ms. **e-f**. H&E staining images of tumor and adjacent normal tissues. Scale bars: 100 μm for Fig. 7b, c, e, f and 1 cm for Fig. 7d.

## Discussion

This study started from the spectral data of the clinically available NIR organic fluorescent dye ICG, providing a more complete depiction of the electronic transition processes among various singlet excited states. We hypothesized and verified that ICG emits visible two-photon fluorescence under 785 nm CW excitation for the first time. On this basis, the conditions required for organic dyes to achieve this luminescence were analyzed, and a new mechanism of CW-excited two-photon fluorescence emission was summarized and proposed. It was subsequently validated across multiple organic dyes of different types. Finally, a CW-excited two-photon fluorescence microscopic system was constructed, achieving three-dimensional optical sectioning imaging of mouse cerebral vasculature and clinical human gastric cancer tissues, thereby validating the capability of this mechanism for high-resolution optical sectioning imaging in highly scattering media.

The core of the new mechanism for CW-excited two-photon fluorescence emission proposed in this study lies in the ESA process of electrons. A key condition for realizing this process is that the real energy level of the intermediate excited state can be populated with a sufficient number of electrons within a short time. Apart from the external factor of excitation power, two intrinsic molecular properties play crucial roles: the lifetime of the intermediate singlet excited state, and the extinction coefficient from the ground state to this intermediate singlet excited state. The former primarily relies on accumulative effects over a certain period to achieve a large population, whereas the latter directly populates a sufficient number of molecules in an extremely short time. Therefore, the relationship among the number of electrons N_1_ in the intermediate singlet excited state, its lifetime τ_1_, the molar extinction coefficient ε_0_ from the ground state to this excited state, and the excitation power P can be expressed as follows:

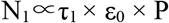

Owing to the new mechanism proposed in this study, current fluorophore systems capable of CW-excited, ESA-based two-photon fluorescence emission can be divided into two categories: the organic dyes systematically explored in this work, and rare-earth-doped nanoparticles (UCNPs)^10^. Organic dyes exhibit a large ε_0_, reaching approximately 10^5^ L·mol^−1^·cm^−1^, but a short τ_1_, typically on the order of hundreds of picoseconds or nanoseconds. Interestingly, UCNPs show the opposite characteristics, with ε_0_ typically as low as approximately 10^2^ L·mol^−1^·cm^−1^, but τ_1_ as long as milliseconds. Therefore, we speculate that if an organic fluorescent dye satisfying the conditions described herein also possesses a longer τ_1_, it would further enhance the two-photon absorption efficiency, potentially enabling two-photon fluorescence emission under even lower-power CW excitation, or even with only LED or sunlight illumination. This could further improve safety from the perspective of the excitation source and promote the application of organic dyes in clinical imaging, infrared sensing, and related fields.

The new mechanism and the constructed method proposed in this study offer multiple advantages. At the fluorophore level, there are currently two main types of fluorophores capable of CW-excited two-photon fluorescence: UCNPs and triplet–triplet annihilation (TTA)-based organic molecules^11–14^. Among these, UCNPs are inorganic systems containing heavy metals, and their in vivo biosafety and stability require further evaluation. They also involve complex synthesis and are difficult to reliably produce at large scale with high consistency^10^. TTA-based organic molecules require the pairing of sensitizers and annihilators, making the synthesis process equally cumbersome. Moreover, owing to the involvement of triplet states, they are highly sensitive to oxygen and exhibit poor in vivo stability^15^. In contrast, the new mechanism proposed herein is entirely based on intramolecular electronic transitions among singlet states in organic molecules, offering good in vivo stability. Moreover, organic dyes such as ICG, OTL38, and SPN-01 possess excellent biocompatibility and high clinical application value. At the imaging system level, we have successfully replaced femtosecond laser sources with much lower-cost CW sources with significantly lower peak power for two-photon excitation. We achieved three-dimensional high-resolution visualization of mouse cerebral vasculature and human tumor tissues, thereby providing detailed and rich information for physician diagnosis and treatment using a low-cost approach. By combining the safer CW excitation source with clinically available organic fluorescent dyes, this study offers an exciting new high-resolution optical sectioning imaging modality for fundamental biological research and clinical medicine.

It is important to note that this new mechanism was discovered and summarized in some classical NIR organic fluorescent dyes. ICG, OTL38, and SPN-01 all belong to the family of NIR organic cyanine fluorescent dyes. These dyes can emit fluorescence in the first near-infrared window (NIR-I) and even NIR-II. Because this class of NIR fluorescent dyes represents a well-established system and encompasses a rich variety of dyes, we have good reason to believe that a large number of organic fluorescent dyes exhibiting this luminescent mechanism will be uncovered as research progresses. In other words, a NIR-I or NIR-II organic fluorescent dye may simultaneously possess frequency downconversion one-photon fluorescence emission and frequency upconversion two-photon fluorescence emission. This bears a strong resemblance to the well-established UCNPs system, while forming a complete and self-contained organic system of its own. The discovery of this new physical mechanism of fluorescence emission is expected to play a revolutionary guiding role in the field of organic fluorescent probe synthesis.

## Materials

ICG, IR-820 and DMSO were purchased from Shanghai Aladdin Bio-Chem Technology Co., LTD. OTL38 was purchased from MedChemExpress (MCE). The aqueous solution of SPN-01 was provided by Professor Zhifei Dai’s group in Peking University. The aqueous solution of Y6CT-NPs was provided by Professor Jianguo Wang’s group in Inner Mongolia University.

## Data availability

The raw data will be made available upon request to the corresponding author. The remaining data are available within the Article.

## Acknowledgements

This work was supported by the National Key R&D Program of China (2024YFF1206700), the National Natural Science Foundation of China (U23A20487), the Dr. Li Dak Sum & Yip Yio Chin Development Fund for Regenerative Medicine, Zhejiang University, China, the Hangzhou Chengxi Sci-tech Innovation Corridor Management Committee funded project.

## Author contributions

J.Q. conceived the idea. S.W. led the experiment with supervision from J.Q.. S.W. and J.Q. discussed the experimental ideas. X.F. collected the human gastric cancer and adjacent normal tissue samples. X.L. performed the H&E staining of the tissues. D.F. carried out the tissue incubation and washing procedures. X.M. conducted the transient absorption measurements. Z.F. provided valuable suggestions for the writing of the manuscript. S.W. processed and analyzed experiment data. W.H. and J.Q. manage the project. S.W. and J.Q. wrote and revised the paper.

## References

1. Helmchen, F. & Denk, W. Deep tissue two-photon microscopy. Nat Methods 2, 932–940 (2005).

2. Hayazawa, N., Tarun, A., Taguchi, A. & Kawata, S. Development of Tip-Enhanced Near-Field Optical Spectroscopy and Microscopy. Jpn. J. Appl. Phys. 48, 08JA02 (2009).

3. Christensen, D. J. & Nedergaard, M. Two-photon in vivo imaging of cells. Pediatr Nephrol 26, 1483–1489 (2011).

4. Cahalan, M. D., Parker, I., Wei, S. H. & Miller, M. J. Two-photon tissue imaging: seeing the immune system in a fresh light. Nat Rev Immunol 2, 872–880 (2002).

5. Cheng, H., Tong, S., Deng, X., Liu, H., Du, Y., He, C., Qiu, P. & Wang, K. Deep-brain 2-photon fluorescence microscopy in vivo excited at the 1700 nm window. Opt. Lett. 44, 4432 (2019).

6. Booth & Hell. Continuous wave excitation two-photon fluorescence microscopy exemplified with the 647-nm ArKr laser line. Journal of Microscopy 190, 298–304 (1998).

7. Hell, S. W., Booth, M., Wilms, S., Schnetter, C. M., Kirsch, A. K., Arndt-Jovin, D. J. & Jovin, T. M. Two-photon near-and far-field fluorescence microscopy with continuous-wave excitation. Opt. Lett. 23, 1238 (1998).

8. Apanasevich, P. A. & Timofeeva, G. I. Resonant Two-Photon Transitions. J Appl Spectrosc 85, 250–254 (2018).

9. Sauter, G., Papapostolou, A., Pollien, A., Boschmann, S., Fuchs, K., Merten, P., Brödner, K., Rominger, F., Freudenberg, J., Bunz, U. H. F., Dreuw, A. & Tegeder, P. Exceptionally High Two-Photon Absorption Cross Sections in Quinoidal Diazaacene-Bithiophene Derivatives. Angew Chem Int Ed 64, e202503073 (2025).

10. Zhou, J., Schuck, P. J. & Jin, D. Upconverting nanoparticles for biomedical applications. Nat Rev Phys 8, 195–207 (2026).

11. Liu, Q., Feng, W., Yang, T., Yi, T. & Li, F. Upconversion luminescence imaging of cells and small animals. Nat Protoc 8, 2033–2044 (2013).

12. Chen, G., Qiu, H., Prasad, P. N. & Chen, X. Upconversion Nanoparticles: Design, Nanochemistry, and Applications in Theranostics. Chem. Rev. 114, 5161–5214 (2014).

13. Zhou, J., Liu, Q., Feng, W., Sun, Y. & Li, F. Upconversion Luminescent Materials: Advances and Applications. Chem. Rev. 115, 395–465 (2015).

14. Magazov, Y., Aliyev, A., Zhumabay, N., Taubaldiyeva, Z., Zhigerbayeva, G. & Nuraje, N. Upconversion materials: a new frontier in solar water-splitting. RSC Adv. 16, 1643–1661 (2026).

15. Huang, L., Kakadiaris, E., Vaneckova, T., Huang, K., Vaculovicova, M. & Han, G. Designing next generation of photon upconversion: Recent advances in organic triplet-triplet annihilation upconversion nanoparticles. Biomaterials 201, 77–86 (2019).

